# A novel murine tail model for radiation-induced skin injury following localized irradiation at varying doses

**DOI:** 10.1101/2025.01.10.632342

**Authors:** Guojian Wang, Na Zhao, Shuang Long, Jining Gao, Qing Zhou, Xinze Ran, Junping Wang, Tao Wang

## Abstract

**Background:** Radiation therapy is one cornerstone of oncologic treatment. Radiation-induced skin injury (RISI) is a dose-limiting complication of radiotherapy. RISI is also common in victims of accidental exposure, which often aggravates the patient’s condition and becomes a difficult medical issue. However, the damage mechanisms of RISI remain unclear, and the prevention and treatment measures are limited. An appropriate animal model holds great significance for addressing these issues.

**Methods:** C57BL/6 mice were employed to establish RISI model by irradiating 2 cm section of the mouse tail with 20Gy, 30Gy, and 40Gy of single irradiation. Skin injuries were scored with a modified semi-quantitative scale. H&E staining, IHC for dopachrome tautomerase (DTC) and Masson staining were used for histopathological evaluations of RISI.

**Results:** We innovatively established animal models of RISI through tail irradiation. The model mice showed typical symptoms of dry desquamation, moist desquamation, ulcers and necrosis in the irradiated area, which were highly similar to the clinical manifestations. Concurrently, we discovered that in the later stage of this model, the interstitial tissue at irradiated site presented a fibrotic phenotype with good dose dependence. It should be noted that tail irradiation causes dynamic changes in skin melanin, with early excessive deposition and late loss, which renders the observation of radiation-induced skin erythema difficult. For this reason, we specifically revised the scoring criteria for RISI.

**Conclusion:** In conclusion, this study established an easily operable and highly reproducible tail irradiation model, providing a novel platform for in-depth research on the mechanisms and translational applications of RISI.

## 1. INTRODUCTION

Radiation-induced skin injury (RISI), also known as radiation dermatitis, is a common complication that occurs in approximately 95% of radiotherapy patients^1-4^. These types of skin injuries can be classified into acute or late (i.e., chronic) categories^1, 2^. Manifestations of acute injuries typically arise within hours to weeks after exposure presenting with symptoms such as erythema, edema, pigment change, depilation, dry/moist desquamation, necrosis, and ulcers as exposure doses increase. Late injuries emerge months to years later leading to the development of delayed ulcers, fibrosis, and telangiectasia. Skin injuries by irradiation can negatively affect the quality of a patient’s life due to pain and premature interruption of radiotherapy, which in turn may compromise disease management^1, 2^. Meanwhile, radiation-induced skin injuries have been documented in numerous nuclear and radiological accidents, significantly impacting the progression and outcomes of radiation injury cases^1-3^. Unfortunately, there are currently no universally accepted standards for the measurement and management of radiation-induced skin injuries. Therefore, the development of effective agents to prevent or mitigate RISI holds significant importance.

When embarking on studies to investigate the injury mechanisms, identify biomarkers for early diagnostic, and screen medical countermeasures (MCMs) for radiation-induced skin injuries, it is essential to select animal models that not only match the proposed action of the approach to be tested but also reflect anticipated human responses^2^. Diverse animal models, including minipigs, guinea pigs, rats and mice, have been established for radiation-induced skin injuries^2, 5-10^. These models offer distinct advantages and limitations necessitating careful selection based on specific research objectives. Due to their skin architectures and thickness closely resembling human skin tissue, larger animal models like minipigs are ideal for study of RISI^2^. However, due to cost considerations, small animal models such as mice are more commonly used. Mice are easier to manipulate genetically and relatively inexpensive. Therefore, cutaneous radiation injury models in mice are often preferred^9-11^. There has been documentation on mouse skin irradiation across different body regions like the dorsal back^10^, legs^12^, head and neck^13^, sole of feet^14^, and ears^11^, yet tail skin irradiation models remain unreported. In this study, a mouse model of tail skin radiation injury has been established through exposure to varying doses. The study found that the clinical manifestations and pathological changes in the mouse tail skin after different doses of irradiation were similar to the typical clinical manifestations and pathological changes of humanskin radiation injury. Given these findings coupled with the unique benefits offered by tail studies, this model holds promise for future research endeavors.

## 2. MATERIALS AND METHODS

### 2.1 Animals

All mice related experiments were approved by Laboratory Animal Welfare and Ethics Committee of the Army Medical University (AMU, Chongqing, China). C57BL/6 male mice (8-10 weeks old) were obtained from SiPeiFu Biotechnology (Beijing, China). Animals were housed under a 12 hours light/dark cycle in a SPF room with water and food provided ad libitum.

### 2.2 Tail irradiation

Before optimizing the development of the tail irradiation model, we initially conducted a comparative histological analysis of the tissues of the mouse tail and dorsal skin. As seen in Fig. 1A, the tail skin consists of well-stratified epidermal cell layers similar to human skin (middle panel for longitudinal section and low panel for transection), while the dorsal skin shows thin epidermal monolayer (up panel). For tail irradiation, the animals were intraperitoneally anesthetized with 1% pentobarbital sodium. Irradiation was administered using an RS2000 device to tail skin of the mice, which were covered by a lead shelter (1 cm in thickness) that exposed a portion of the tail region (starting 2 cm from root of the tail and extending for a length of 2 cm, Fig. 1B and 1C). The mice received a single exposure to 160 kV X-rays of 20, 30, and 40 Gy at a dose rate of 1.265 Gy/min.

**Figure 1.**
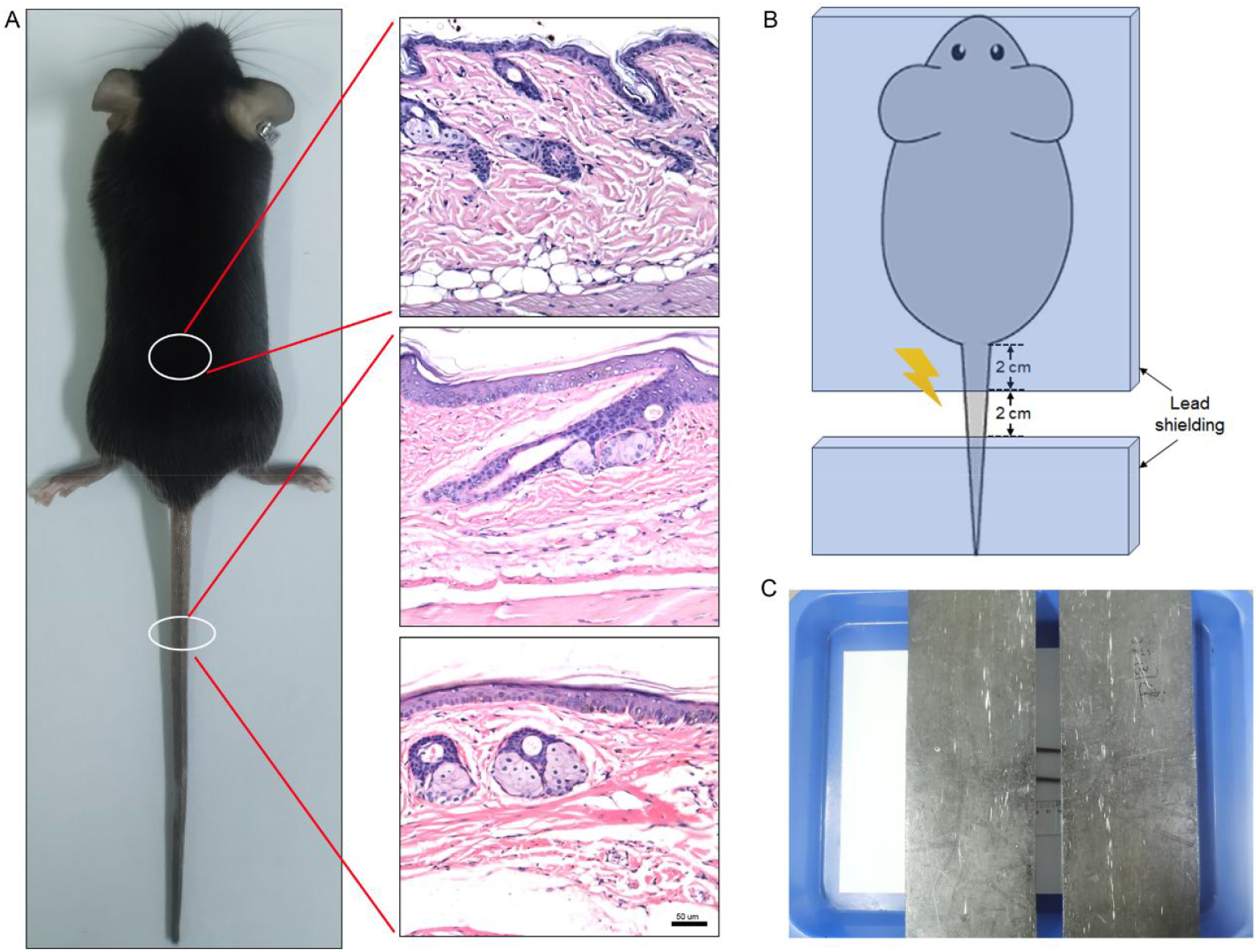
Experimental design. **(A)**, Histological comparison of skin tissue in the back and tail of mice. Left, schematic diagram of sampling sites in mice. Right, up panel: histological section of murine dorsal skin tissue; middle and low panels: longitudinal and cross histological sections of murine tail skin tissue. **(B)**, Illustrative diagram of irradiation for the mouse tail model. (**C)**, Actual scene of irradiation in the mouse tail model.

### 2.3 Score evaluation

The tail skin tissues of C57BL/6 mice naturally contain melanin deposits, resulting in a darker pigmentation^15^. In addition, there are significantly increased melanin deposition in tail skin following irradiation. Thus, the conventional indicator of radiation-induced skin injury in other models, erythema, cannot effectively be used in our new model. Therefore, we have adjusted the criteria for scoring skin injury in this model based on alternative standards^12^ (Table 1 and Fig. 2). Skin damage was measured by two unblinded observers approximately weekly for 8 weeks.

**Table 1.** Score evaluation criteria for radiation induced skin damage.

| Score | Skin changes |
| --- | --- |
| 1.0 | No effect |
| 1.5 | Slight hyperpigmentation, mild dry skin. |
| 2.0 | Moderate hyperpigmentation, dry skin. |
| 2.5 | Marked hyperpigmentation, dry desquamation. |
| 3.0 | Dry desquamation, minimal dry crusting, moderate depigmentation. |
| 3.5 | Dry desquamation, dry crusting, superficial minimal scabbing, severe depigmentation. |
| 4.0 | Patchy moist desquamation, edema, moderate scabbing. |
| 4.5 | Confluent moist desquamation, ulcers, large deep scabs. |
| 5.0 | Open wound, full thickness skin loss. |
| 5.5 | Necrosis. |

**Figure 2.**
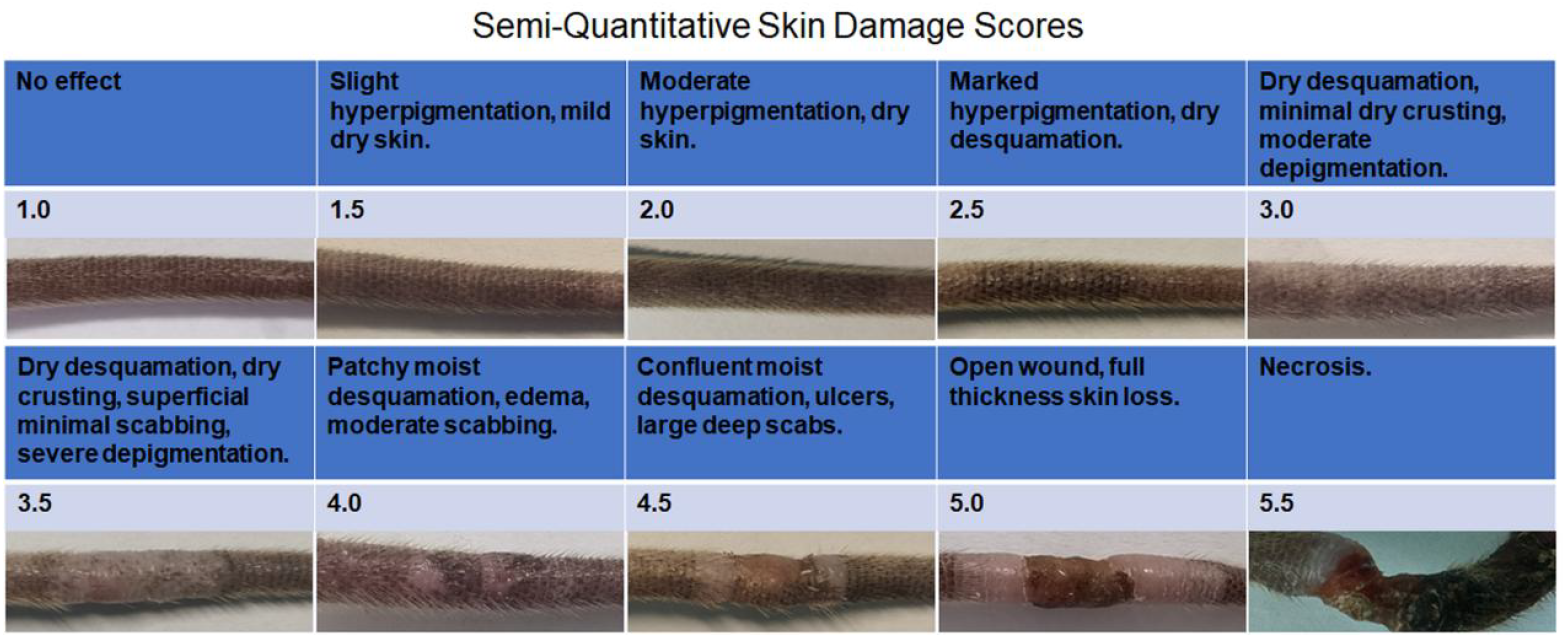
The modified semi-quantitative skin damage scores for radiation-induced skin injury with representative images.

### 2.4 Tissue staining and histopathological analysis

The tail tissues of mice were taken at each time point, fixed in 10% neutral formaldehyde, and subsequently subjected to decalcification for one month. Specimens were then embedded in paraffin, cut into 4-um sections, stained with hematoxylin and eosin (H&E). Masson trichrome staining was employed to evaluate skin fibrosis according to the manufacturer’s protocols. Images of HE and Masson trichrome staining were scanned by the Precipoint M8 Digital Microscope & Slide Scanner (Precipoint GmbH, Germany), and analyzed with the ViewPoint BETA software system. Epidermal thickness was measured at ten different positions per sample and averaged to one data set^16^. Epidermal basal cell density was determined by total basal cells per mm of basal membrane^17^.

### 2.5 Immunohistochemistry

Immunohistochemistry (IHC) staining in skin tissues were performed as previously described. Briefly, tissue sections were stained with rabbit anti-DCT (1:200, Abcam). Then, DAB chromogenic system was used for final chromogen. Images were collected by the Precipoint M8 Digital Microscope & Slide Scanner. Epidermal DCT positive cells were quantitified as the number of positive cells per mm epidermis in the ViewPoint BETA software system.

### 2.6 Statistical analysis

Experimental data were analyzed using GraphPad Prism 6.0 software. Data were shown as mean ± SD. Statistical analysis was performed by

Student’s *t* tests (the *U*-test was performed to test differences between independent groups at different time points during wound healing). A *P* value <0.05 was considered to have statistical significance.

## 3. RESULTS

### 3.1 Score evaluation criteria of semi-quantitative skin damage for tail model

There are many different scoring scales of skin injury for cutaneous radiation injury in both clinical practices and preclinical animal studies^2^. Among the available visual scoring scales, the Kumar scale is widely used in various preclinical animal models such as mice, rats and guinea pigs^7, 8, 12^. The Kumar scale, originally developed for assessment of radiation injury in the hind leg of mice, ranges from 1.0 (no effect) to 5.5 (necrosis) based on erythema, desquamation (dry and moist), ulcers and necrosis^12^. For the convenience of evaluating the presence and degree of erythema, the Kumar scale is suitable for animals with light-colored skin or areas with less pigmentation. The skin of the mouse tail is rich in melanocytes, featuring obvious pigment deposition and a dull color^15^. Meanwhile, ionizing radiation can cause premature differentiation of melanocyte stem cells, further darkening the skin color^18^.

These factors make it impossible to use the erythema index as a measure of radiation damage to the tail skin. During this research, we discovered that the degree of melanin deposition and loss in the irradiated region was closely associated with the severity of skin radiation injury. Accordingly, we incorporated the changes in melanin as one surrogate indicator for erythema into the new scoring scale (Table 1). Furthermore, representative macroscopic observation images corresponding to various scores of the scale have been selected for the facilitation of application (Fig. 2).

### 3.2 Radiation induces dry desquamation with dynamic hyperpigmentation and hypopigmentation in murine tails after single 20 Gy irradiation

Irradiation of the murine tails was executed at varying doses, and the skin injury reactions were observed and analyzed. Our findings indicate that different irradiation doses result in distinct characteristic changes in skin injuries. When a single dose of 20 Gy was administered to the tail skin, the primary alteration was dry desquamation, accompanied by dynamic fluctuations in melanin deposition and loss. As depicted in Fig. 3A, the irradiated area progressively became dry, leading to pronounced dry desquamation and hair loss. On the other hand, the increase in melanin deposition became notably apparent within two weeks post-irradiation, peaking at week four before gradually diminishing thereafter. And, the state of hypopigmentation in the irradiated area of the tail skin can persist for more than 14 weeks. Then, skin injury was scored according to the modified scale (Table 1). Our findings indicated that the skin damage score progressively escalated in conjunction with the advancement of dry desquamation, while no instances of moist desquamation were observed throughout the entire process (Fig. 3B). Further histological observations revealed that epidermis of the tail skin tissue was significantly thicker at 8 weeks after 20 Gy irradiation, with a marked increase in the thickness of the stratum corneum, which was consistent with the phenotype of dry desquamation (Fig. 3C).

**Figure 3.**
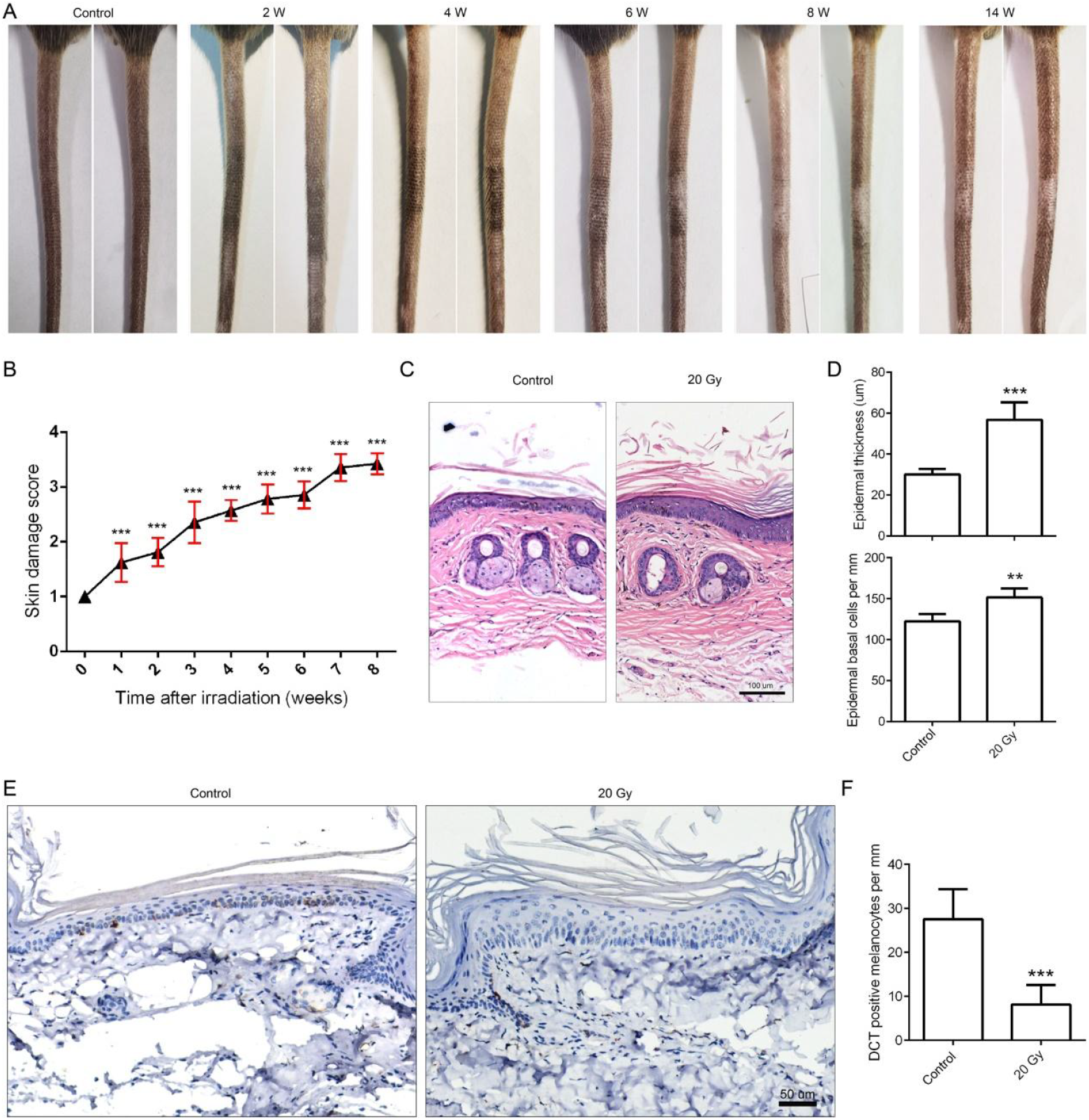
Radiation induces dry desquamation in the tails of mice following a single exposure to 20 Gy. **(A)**, Representative images of mouse tails at various time points following localized irradiation with 20 Gy. (**B)**, Skin injury was scored according to the modified skin damage scoring standards. (**C)**, Histological analysis of mouse tail skin tissues in both control and 20 Gy irradiated mice following 8 weeks post-irradiation. (**D)**, Quantitative analysis of the epidermal thickness (up panel) and basal cell density (low panel) of the tail skin tissues. (**E)**, Representative images of IHC staining for DCT in mouse tail tissues. (**F)**, Quantitative analysis of the DCT-positive epidermal keratinocytes. ^**^ *p*<0.01, and ^***^ *p*<0.001 versus nontreated control mice.

Histopathological semi-quantitative analysis demonstrated that both the epidermal thickness and density of basal cells at 8 weeks after 20 Gy irradiation were significantly elevated compared to those of the control tail tissue (Fig. 3D). To further evaluate the damage status of melanocytes, we conducted dopachrome tautomerase (DCT) immunohistochemical staining on the tail skin at 8 weeks post-irradiation. The results indicated that almost no DCT-positive cells were detectable in the irradiated group, and their quantity was significantly decreased (Fig. 3E and 3F).

### 3.3 Radiation induces moist desquamation of skin in murine tails after single 30 Gy irradiation

As seen in Fig. 4A, a single 30 Gy irradiation to the mouse tail leads to typical moist desquamation. The skin in the irradiated area presented marked moist desquamation in the fifth week, and superficial ulcers formed in the sixth week. Subsequently, it gradually recovered, and the wound was nearly completely healed by the eighth week. Upon observation of the entire wound healing process, it was found that all wounds healed after 30Gy irradiation, without forming chronic ulcers. Furthermore, the scoring results also showed that the skin injury caused by the 30Gy dose reached its peak at week 6, and then decreased (Fig. 4B).

**Figure 4.**
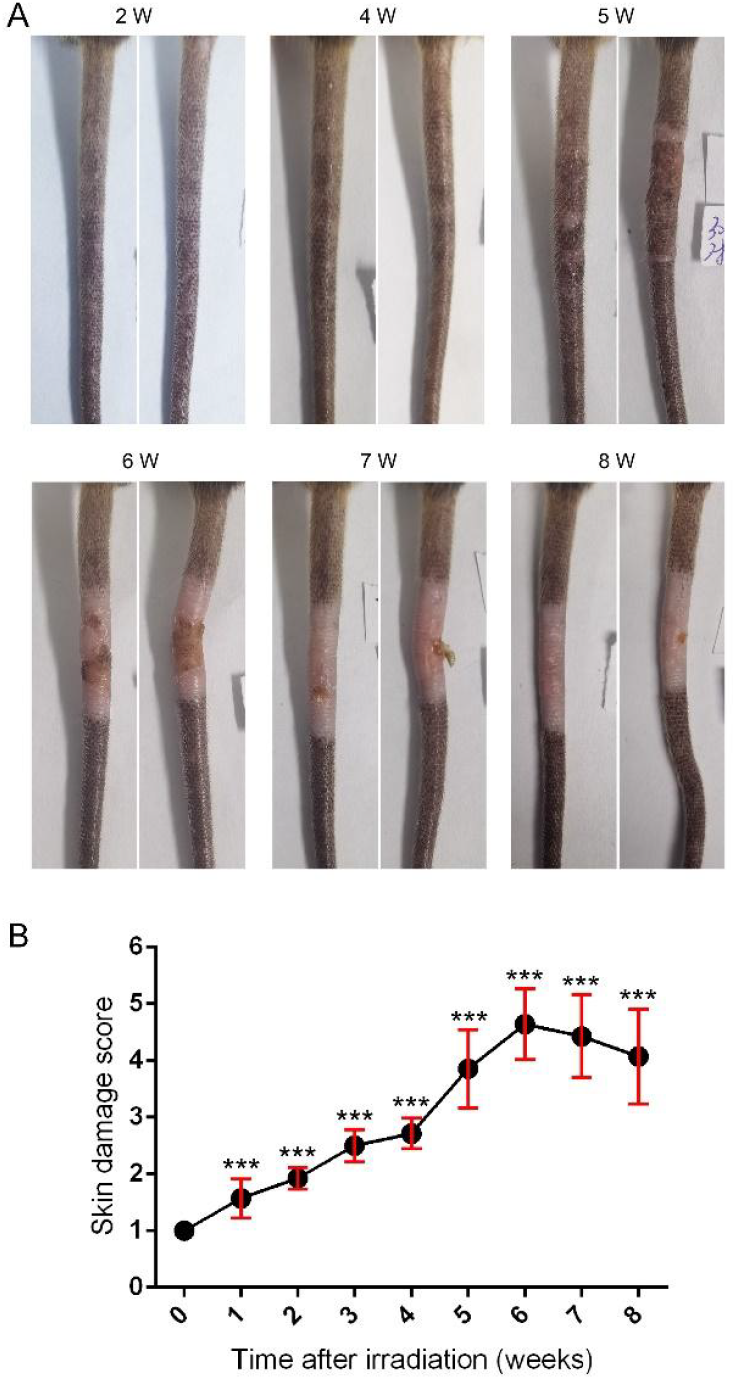
Radiation IR elicits moist desquamation in the tails of mice subsequent to a single exposure to 30 Gy. **(A)**, Representative images of mouse tails at various time points following localized irradiation with 30 Gy. (**B)**, Skin injury was scored according to the modified skin damage scoring standards.

### 3.4 Radiation induces non-healing ulcers in murine tails after single 40 Gy irradiation

When the irradiation dose was raised to 40 Gy, the mouse tail mainly presented with chronic radiation ulcers. As shown in Fig. 5A, the irradiated skin exhibited obvious ulcerative alterations as early as the fifth week, and the subsequent repair progressed slowly. Only a small portion of the ulcers could completely heal, but the majority remained unhealed by the eighth week. The results of the skin damage scoring were in accordance with the gross observations, reaching a plateau at the fifth week after irradiation and persisting until the observation endpoint at the eighth week (Fig. 5B). Further histological observations and analysis revealed that the tail skin showed marked thinning of the epidermis, disappearance of sebaceous glands, and atrophy of hair follicles in the third week after irradiation. By the fourth week, there was a clear gap between the epidermis and dermis, suggesting a severe disruption of their connection. By the eighth week, the tail skin that had healed well showed a markedly thickened epidermis and a well-filled interstitial tissue (Fig. 5C and 5D).

**Figure 5.**
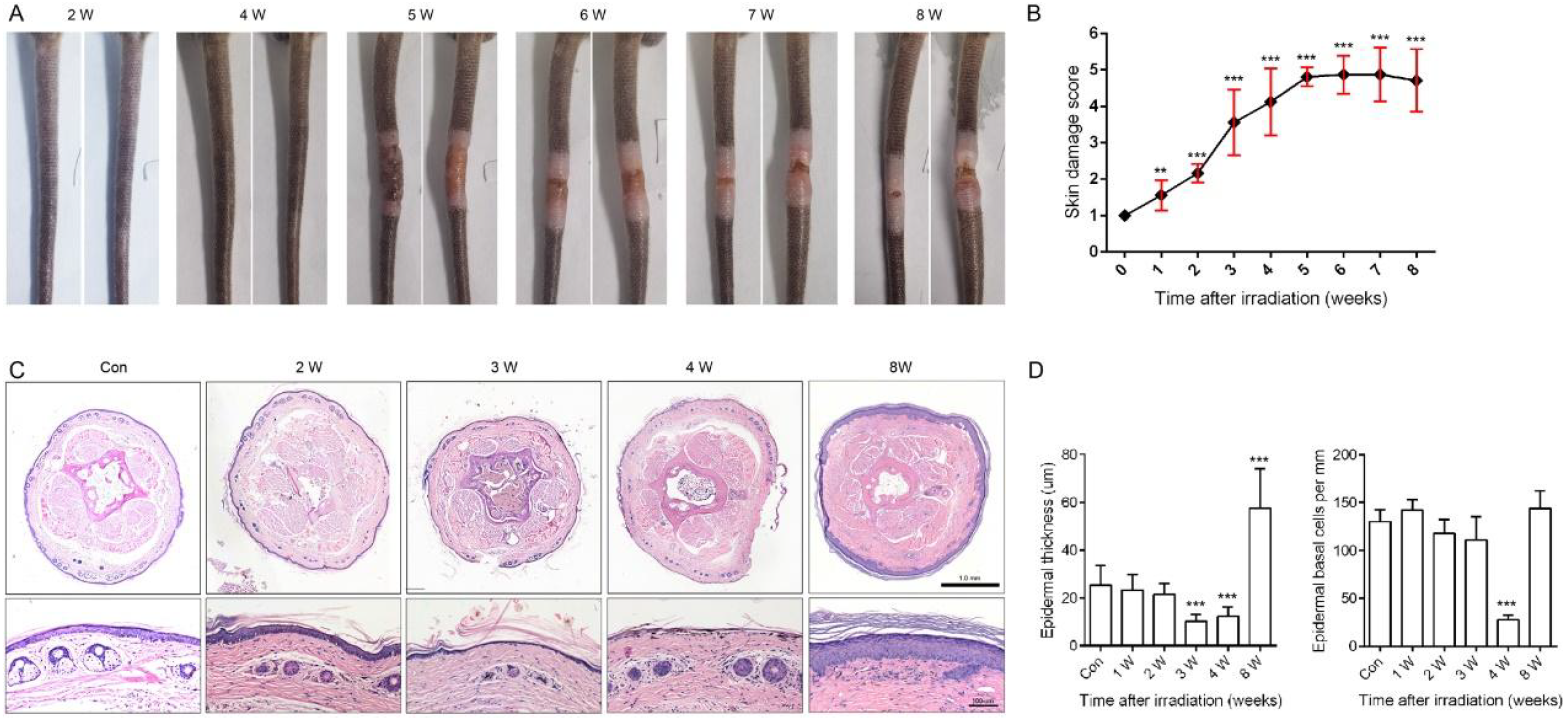
Dynamic observation of the injury manifestations and histopathological changes in the tails of mice after irradiation with 40 Gy. **(A)**, Representative images of mouse tails at various time points following localized irradiation with 40 Gy. (**B)**, Skin injury was scored according to the modified skin damage scoring standards. (**C)**, Representative images of H&E staining for mouse tails at various time points following localized irradiation with 40 Gy. (**D)**, Quantitative analysis of the epidermal thickness (left panel) and basal cell density (right panel) of the tail skin tissues

As previously mentioned, at the eighth week post-irradiation with 40 Gy, a small portion of ulcers healed completely (Fig. 6A, left panel), but most injuries remained unhealed, presenting as significant swelling due to exudate scabbing and drying contraction at the wound site (Fig. 6A, middle panel), which could lead to visible necrosis if not effectively alleviated (Fig. 6A, right panel). Pathological observations of the cross-section of the mouse tail tissue disclosed that, in contrast to the normally healed mouse tail tissue (Fig. 6B, up panel), the unhealed tail exhibited severe edema caused by the dry scab, histologically presenting as swelling and the occurrence of obvious large necrotic areas (Fig. 6B, low panel). Further histological observations of the longitudinal section of the ulcer revealed that the ulcer site contracted conspicuously, the epidermal layer at the ulcer margin was significantly thickened, and the entire ulcer was accompanied by extensive infiltration of inflammatory cells (Fig. 6C).

**Figure 6.**
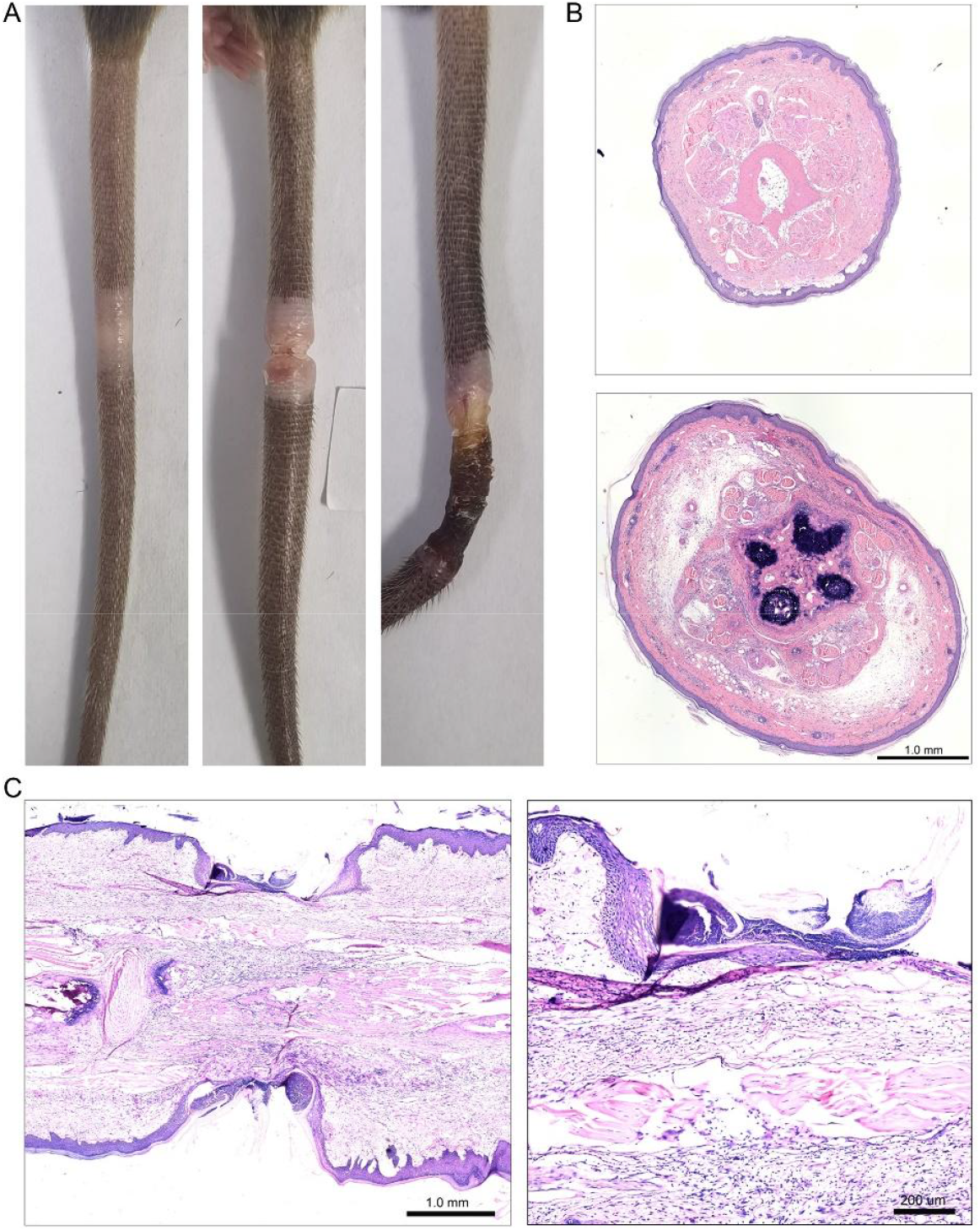
Radiation induces ulcers and necrosis in the tails of mice following a single exposure to 40 Gy. **(A)**, Representative images of the three outcomes of the mouse tail at 8 weeks post-40Gy irradiation, healed (left panel), ulcer (middle panel), and necrosis (right panel). (**B)**, H&E staining for cross sections of mouse tails from healed (up panel) and ulcer (low panel) samples. (**C)**, A longitudinal sectional observation of the non-healing ulcer in the mouse tail, with the locally enlarged image on the right panel.

### 3.5 Radiation induces dermal fibrosis in murine tails

Fibrosis represents a significant late-effect manifestation resulting from ionizing radiation exposure^19^. Pathological examination of H&E sections revealed that as the irradiation dose increased, collagen fibers in the dermis exhibited marked thickening, accompanied by a substantial reduction in the area of blank spaces observed in cross-sectional views (Fig. 7A). Additionally, Masson staining demonstrated that post-irradiation, dermal collagen in the mouse tail not only became thicker but also more abundant, with an increasingly pronounced fibrotic phenotype correlating with higher irradiation doses (Fig. 7B).

**Figure 7.**
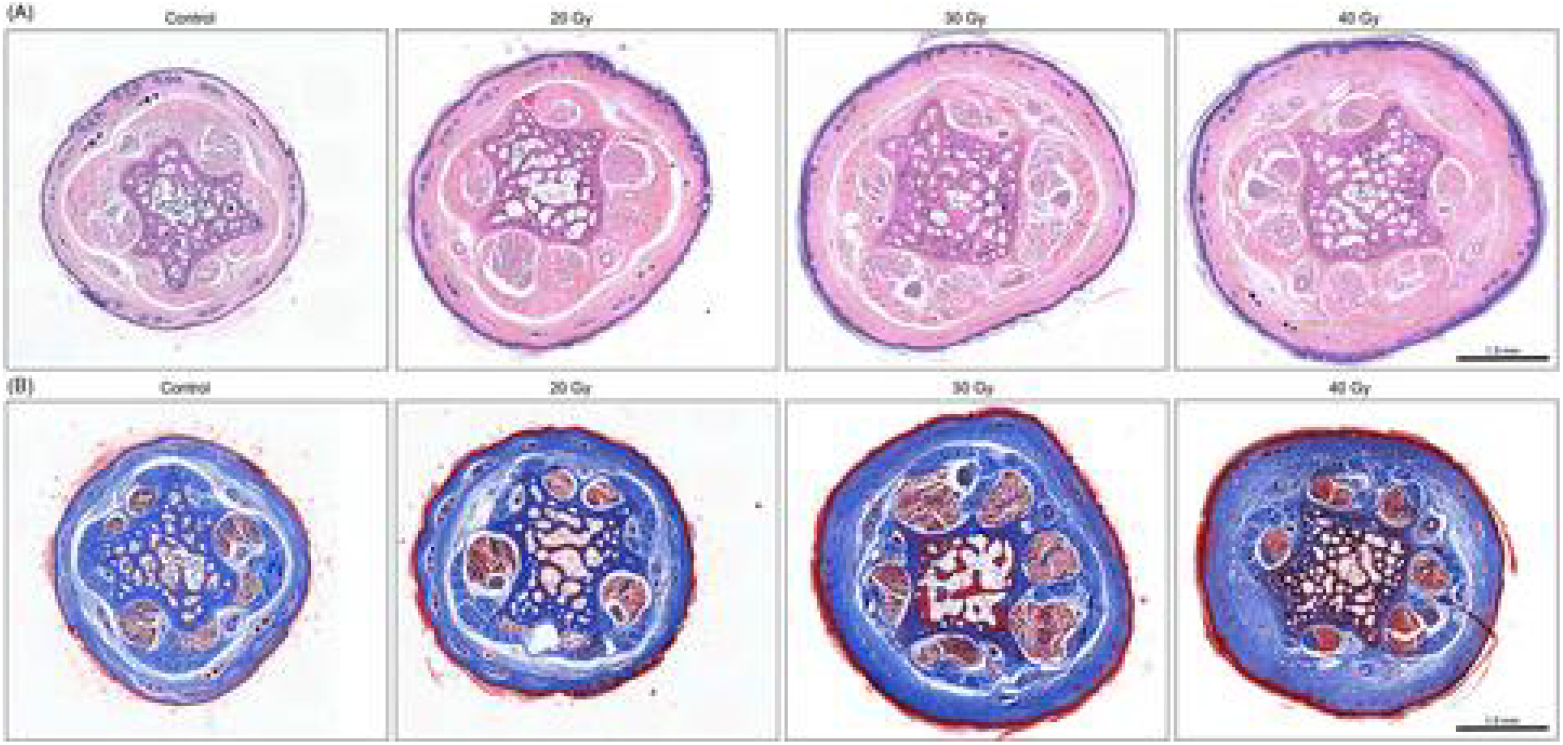
Radiation induces dermal fibrosis in murine tails. **(A)**, Representative images of H&E staining for cross sections of mouse tails after 8 weeks post-irradiation. (**B)**, Representative images of Masson staining for cross sections of mouse tails after 8 weeks post-irradiation.

## 4. DISCUSSION

The successful development of novel medical countermeasures against radiation injuries requires the development of well-characterized animal models which clinically reflect anticipated injury patterns and responses of human^2^. In the current work, we have developed a novel model of radiation-induced skin injuries in mice, characterized by typical manifestations including dry desquamation, moist desquamation, ulcers, and fibrosis following exposure to various doses of localized radiation in the tail region. We propose that this model will facilitate the development of MCMs for radiation-induced skin injuries.

The development of animal models to simulate human radiation-induced skin injuries is a crucial step in elucidating the injury mechanisms, thereby facilitating the discovery of MCMs^2^. It is commonly held that both anatomical and physiological similarity in skin tissues between humans and the animal species should be given priority when selecting model animals for studying radiation-induced skin injuries^2, 5, 6^. The anatomy and physiology of large pig models seem to most closely resemble that of humans, with similar skin thickness, hair and sweat glands. However, its high cost and demanding house conditions restrict its wide application. In addition, both mini pig and guinea pig models, which have a high degree of similarity in anatomy and physiology to human skin, are good alternatives^5, 7^. Mice are successful model organism widely employed in the field of radiation medicine research, and their utilization in the domain of cutaneous radiation injury studies is also considerable. The primary reasons for this are that mice are easy to breed, handle, and genetically manipulate. The skin of mice at different body parts exhibits some degree of heterogeneity. The skin in most regions of mice, such as the trunk, neck, and limbs, is loose and highly mobile, featuring thin epidermis with sparse basal cells. Once injured, wound contraction makes a considerable contribution to the healing process^3, 20^. Although both histological and repair characteristics are quite distinct from those of human skin, radiation injury models of mouse skin on the dorsal back and legs have been frequently reported^10, 12^. The skin at special sites such as the ears, feet, and tails of mice consists of well-stratified epidermal cell layers similar to human skin^21^. Previous studies have reported on cutaneous radiation injury models on the ears and feet^11, 14^, but no model for the tail has been developed. And this study made useful explorations in this area.

We contend that the mouse tail skin radiation injury model has obvious advantages over other types of mouse models. At the operational level of model establishment, this model is easier to achieve effective shielding of other parts of the body, making the model more convenient. In term of effects, this model demonstrates favorable dose-response characteristics, presenting typical phenotypes such as dry desquamation, moist desquamation, ulceration, necrosis, and fibrosis with increasing doses. The high degree of uniformity of effects within each dose group indicates that this approach has excellent reproducibility. Furthermore, from the perspective of drug screening, this model facilitates a more effective evaluation of localized MCMs administration interventions, while also providing a clearer and more accessible assessment of phenotypic characteristics. Of course, there are also some limitations to the mouse tail model of cutaneous radiation injury.

One particularly notable issue is that the existing scoring standards cannot be completely transplanted for the assessment of mouse tail model. Mouse tail skin has a relatively intense melanin deposition. Moreover, the premature differentiation of melanocyte stem cells after irradiation further intensifies the melanin deposition^18^. This makes it challenging to observe radiation-induced erythema damage and, consequently, it is difficult to conduct a general assessment of the damage using traditional scoring scales^22^. Therefore, we have made revisions to Kumar scoring system, replacing the evaluation related to erythema with the deposition and loss of pigmentation after irradiation. Preliminarily, it is capable of reflecting the early skin damage situation. Subsequently, we will optimize the conditions and evaluation standards of the mouse tail irradiation model in albino mice such as Babl/c.

One prominent characteristic of the mouse tail irradiation model is the dynamic early hyperpigmentation and subsequent hypopigmentation in irradiated skin, which is seldom mentioned in other animal models. This is mainly due to the fact that the mouse tail skin of C57BL/6 mice used in our study contains a rich supply of melanocytes^15^. Previous research has discovered that ionizing radiation-induced DNA damage can lead to the depletion of melanocyte stem cells by inducing them differentiating into mature melanocytes^18^. The dynamic changes of melanin in the irradiated area that we observed are essentially in line with this report, which also indicates that this model may become a novel method for researching the renewal and differentiation of skin melanocyte stem cells.

One of the outstanding advantages of this model is that it presents a good dose-response relationship, with typical phenotypes such as dry desquamation, moist desquamation, ulceration, and necrosis appearing on the tail of mice after different doses of irradiation. Therefore, these models will provide convenience for the screening and identification of more targeted and precise MCMs.

It should be noted that the results and conclusions presented in this paper are mainly based on the experimental studies of single-dose irradiation, while clinical cutaneous radiation injuries caused by radiotherapy are often the results of fractionated irradiation^23-25^. Therefore, it is necessary to further explore the mouse tail skin radiation injury model of fractionated irradiation in the future.

## 5. CONCLUSION

In summary, we established a novel animal model that may serve as a basis for further radiation research on skin injury. Our research results demonstrate that administering different doses of irradiation to the tails of mice can present a favorable dose-effect relationship, manifesting typical phenotypes such as dry desquamation, moist desquamation, ulcers, and necrosis. This model offers a novel platform for investigating the mechanisms of skin injury caused by ionizing radiation and the screening of preventive and therapeutic strategies for these clinical issues.

## AUTHOR CONTRIBUTIONS

T.W. conceptualized the study and designed the experiments; G.W., N.Z., S.L., and Q.Z. performed the experiments; T.W., J.N., X.Z., and J.W. reviewed the literature and analyzed data; G.W., and N.Z. wrote the initial manuscript; T.W. revised the manuscript. All authors reviewed and approved the final version of the manuscript.

## ACKNOWLEDGEMENTS

This work was supported by the Chinese National Natural Science Foundation (Grant No. 82172219).

## DATA AVAILABILITY STATEMENT

The data that support the findings of this study are available from the corresponding author upon reasonable request.

